# Carcinogen-induced preinvasive human squamous lung cancer is characterised by a novel atypical basal cell state and presents opportunities for chemoprevention

**DOI:** 10.64898/2026.09.17.748423

**Authors:** Wenrui Guo, Holly A. R. Giles, Alice Chernaik, Yasin Memari, Joanna Xie, Daniella Black, Anne Hilgendorff, Christina Gabriel, Onder Yildirim, Helen Davies, Martin Goddard, Lauren D’Sa, Otmar Schmid, Serena Nik-Zainal, Namshik Han, Frank McCaughan

## Abstract

There are no adequate carcinogen-induced models of the earliest stages of human squamous lung cancer (LUSC), limiting the development of effective chemoprevention strategies. We developed a novel human model of early LUSC by exposing differentiated primary human airway cells to a chemical carcinogen known to cause murine LUSC. Carcinogen exposure induced a phenotype recapitulating the earliest stages of human LUSC and associated with the emergence of a novel population of atypical basal cells (ABCs). ABCs express key cancer hallmark markers but without evidence of either common LUSC-associated mutations or an emergent major clonal/subclonal population. We established the clinical relevance of this model by demonstrating enrichment of the ABC expression profile in independent clinical precancer specimens. We then show that capivasertib, a clinically approved kinase inhibitor, prevented the emergence of the abnormal phenotype and the accumulation of cells with an active DNA damage response. Further, capivasertib reversed the established phenotype; and remained effective when nebulised *in vitro*, mimicking therapeutic inhalation. Together, these findings establish a tractable human model of carcinogen-induced early LUSC and provide a proof-of-principle for repurposing targeted therapies for the prevention of lung cancer.

## Introduction

Lung cancer is responsible for the largest proportion of cancer-related deaths^1^. Squamous lung cancer (LUSC) is responsible for approximately 30% of lung cancers and is a devastating disease for which there are no approved small molecule inhibitors. A minority of LUSC patients have a durable response to immunotherapy^2^, but there remains a significant unmet clinical need to improve outcomes.

In addition to smoking cessation and early detection, there is much interest in developing therapeutic strategies for early intervention and/or chemoprevention of lung cancer^3^. There are currently no licensed compounds for these purposes. The prospect of chemoprevention would be significantly improved by using rational *in vitro* (human cell) models of early LUSC that mimic the clinical disease and can be used to test potential prevention strategies. However, this has been challenging - the most commonly used cell lines for LUSC poorly recapitulate the disease and do not express the canonical markers of LUSC used in the clinic^4^. Further, LUSC-derived patient-derived LUSC organoids are challenging to develop despite occasional positive reports^5^.

Carcinogen-exposed murine models reflect the complexity of epithelial carcinogenesis and therefore have some advantages over transgenic models^6^. One of the most useful LUSC models is a carcinogen-exposed murine model in which animals are exposed over 12-52 weeks to N-nitroso-tri-chloroethylurea (NTCU), leading to preinvasive and then invasive LUSC^7^. It was first reported more than three decades ago and has previously been used for chemoprevention studies^8^ and to study basal cell dynamics in early murine LUSC^9^. However, the significant toxicity, variable penetrance and extended time course of carcinogen exposure present significant challenges to its widespread utility. Further, there are well-documented issues in translating findings from mouse models to the clinic^10^. There is therefore an unmet need for rational carcinogen-based human cell exposure models.

Here we exposed long-term differentiated adult human airway cells at the air-liquid interface (ALI) to NTCU. We demonstrate the emergence of a phenotype that reflects the earliest stages of human squamous carcinogenesis and describe an atypical basal cell (ABC) population with evidence of both plasticity and cancer hallmarks. We then repurpose a licensed pan-AKT inhibitor, capivasertib, and demonstrate its efficacy not only in preventing the emergence of the early LUSC phenotype, but in reversing the phenotype once established. We have established a novel human model of the earliest stages of lung squamous carcinogenesis with broad implications for our understanding of the epithelial response to carcinogens and demonstrate a potential to intervene successfully at the earliest stages of LUSC development.

## Results

### Differentiated Human Bronchial Epithelial Cells (HBECs) develop a squamous lung precancer phenotype in response to the carcinogen NTCU

We adapted the murine model of carcinogen-induced LUSC^7^ to a short-term human carcinogen challenge model using primary patient-derived human ALI cultures. The ALI model is a stable long-term culture that recapitulates key features of the adult human airway epithelium similar to those described using the term “epithelioids”^11^.

KRT5^+^/TP63^+^ human airway basal cells are the stem cells of the adult human airway. Basal cells were expanded *ex vivo* from bronchial brushings (**Figure 1A and 1B**) and differentiated into typical airway pseudostratified epithelia on transwells at ALI. Basal cells were cultured on transwell inserts with or without NTCU, with bi-weekly media changes before histological, immunohistochemical or single-cell transcriptomic analyses (**Figure 1C**).

**Figure 1.**
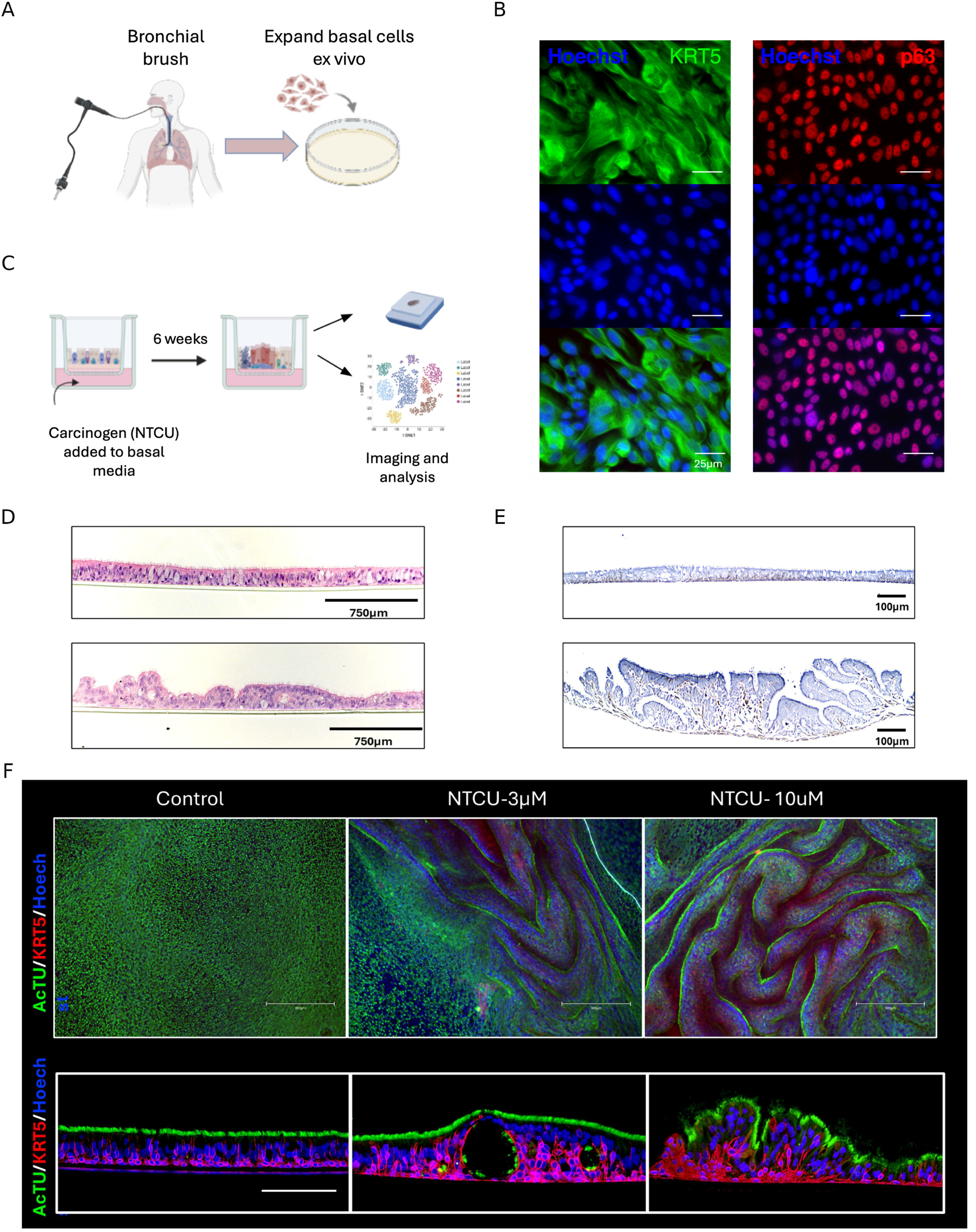
Establishing an *in vitro* model of early lung squamous carcinogenesis. (A) Schematic overview of the experimental workflow. Human airway basal cells were isolated from bronchial brushings and expanded *ex vivo* in standard two-dimensional culture. (B) Immunofluorescence (IF) staining of expanded basal cells for the canonical basal cell markers KRT5 (green) and p63 (red), with nuclei counterstained with Hoechst (blue). Scale bar, 25µm. Representative of four biological replicates. (C) Schematic representation of the air–liquid interface (ALI) culture system used for carcinogen exposure. Differentiated airway epithelial cultures were maintained at ALI and treated with 0-30µM N-nitroso-tri-chloroethylurea (NTCU) for six weeks. (D) Representative hematoxylin and eosin (H&E) staining showing a normal pseudostratified epithelium in control cultures and an abnormal phenotype characterised by epithelial thickening and dysplastic cells in NTCU-treated cultures (10µM). Scale bar, 750µm. Representative images from more than five biological replicates. (E) Immunohistochemistry (IHC) for KRT5. In control cultures (top) the KRT5 staining is generally restricted to the cells directly on the transwell membrane. After six weeks of 30µM NTCU exposure (bottom) the dysplastic phenotype is associated with a thickening of the epithelium and expansion of KRT5^+^ cells towards the putative lumen. Representative images from two biological replicates at the 30µM dose. Scale bar, 100µm. (F) IF staining for acetylated tubulin (green) and KRT5 (red), with nuclei counterstained with Hoechst (blue), demonstrating dose-dependent disruption of epithelial organisation and KRT5^+^ basal cell expansion following NTCU treatment. Scale bar, 300µm (top) and 150µm (bottom). Representative images from three biological replicates.

After exposure to NTCU, marked phenotypic changes are observed (**Figure 1D**). These include a dose-dependent increase in cellularity with formation of pseudopapillae. There is hyperplasia with focal thickening of the epithelial layer suggestive of loss of growth inhibition. The thickening is associated with an expansion of KRT5^+^ basal cells particularly in focal areas of hyperplasia (**Figures 1E-F**) suggesting that the carcinogen-induced phenotype is associated with an expansion of basal cells. This is consistent with a recent description of an NTCU mouse model in which preinvasive lesions were associated with altered basal cell dynamics^9^.

### Single-cell transcriptomics defines an atypical, basal cell-derived population in response to carcinogen exposure

We characterised the human NTCU model using single cell transcriptomics to identify changes in epithelial cell dynamics that may underlie the observed phenotype. We integrated data from 41,113 cells derived from three individuals exposed to either vehicle control or NTCU at two concentrations (**Supp. Table 1**).

We first confirmed that our ALI epithelioids capture the constituent cell types of the human large airway epithelium. Cells were annotated based on expression of marker genes and cross-referenced with known markers from the published literature^9, 12, 13–17^ (**Supp. Table 2**). The ALI model accurately recapitulated differentiated proximal airway epithelia including ciliated, basal and secretory cells (goblet cells, club cells) as well as rarer populations such as ionocytes and tuft cells (**Figure 2A, Supp. Figure 1A)**. We noted a novel population positioned in the centre of the Uniform Manifold Approximation and Projection (UMAP), between the basal and ciliated clusters which we termed atypical cells.

**Figure 2.**
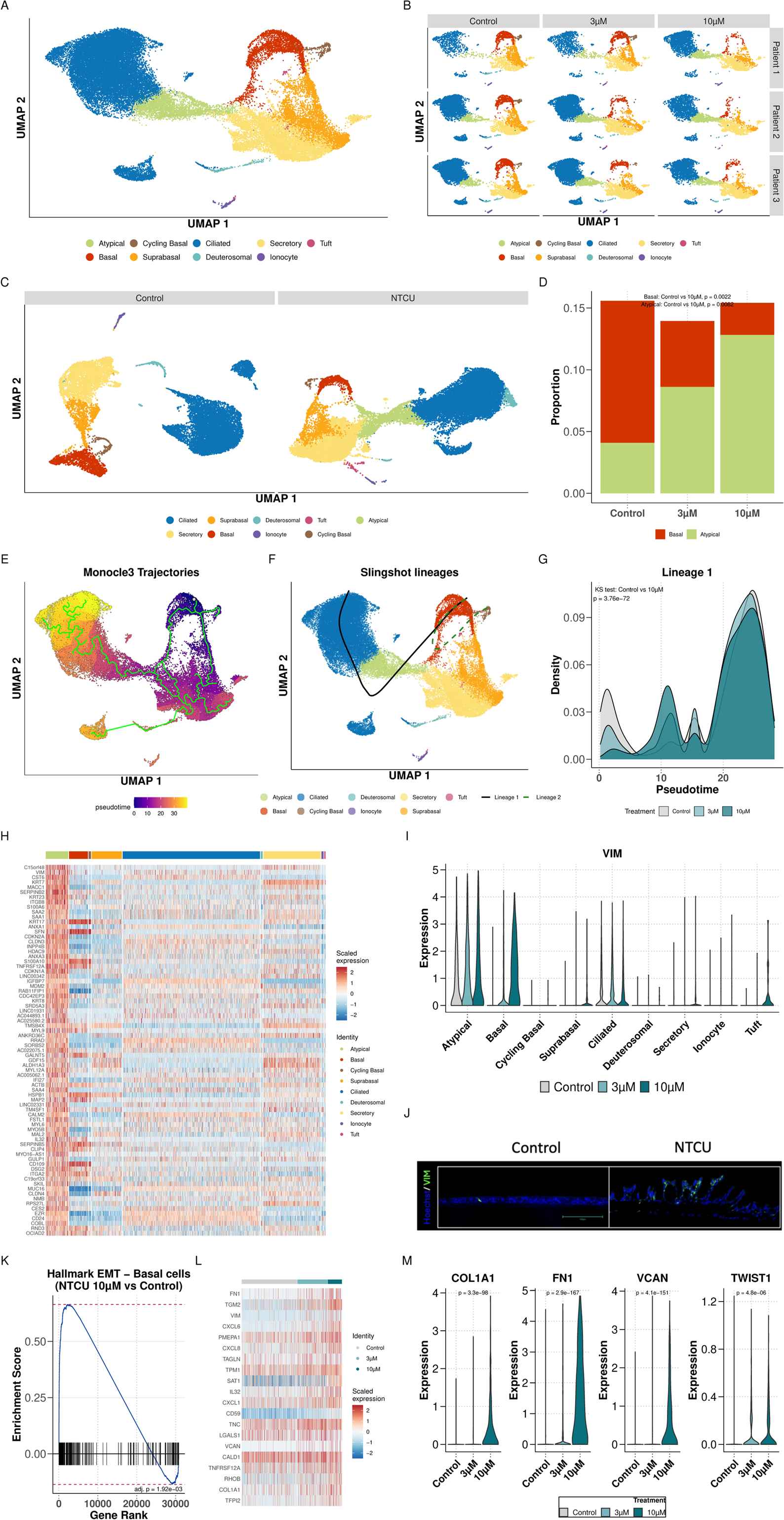
Emergence of atypical cells in model of early lung squamous carcinoma. A) Uniform Manifold Approximation and Projection (UMAP) visualisation of differentiated human airway epithelium cells in ALI cultures. The UMAP represents integrated single cell transcriptomics data from 41,113 cells from nine samples (three patient samples 1, 2 and 3, each treated with Control and NTCU in two concentrations (3µM and 10µM). Cluster colour corresponds to cell type. B) UMAP visualisations of the nine samples, as in (A), split by patient sample (1, 2, 3) and by treatment (Control, 3µM NTCU, 10µM NTCU). C) UMAP visualisations of separately integrated single cell transcriptomics data from (left) the three patient samples treated with vehicle control and (right) the six samples comprising the three patient samples treated with NTCU at two concentrations (3µM and 10µM). D) Bar plot showing relative abundance of basal cells and atypical cells in the control and NTCU-treated groups. Abundance is representative of the three patient sample replicates in each group. Bar colour indicates cell type. Statistical significance of cell-type proportion changes (Control vs 10µM) was assessed using Propeller (Basal *p*=0.0022, Atypical *p*=0.0082). E) Trajectory analysis using Monocle3 to investigate epithelial cell dynamics corresponding to the changes in cell composition seen in (D). Analysis was run on all cell types shown in (A), cells were ordered with the basal population set as starting node (grey circle), the green lines depict possible trajectories, scale bar depicts pseudotime. F) Cell lineage inference for basal, suprabasal, atypical and ciliated cell populations, produced with Slingshot. Lineages are projected onto complete UMAP. Lineage 1 (black solid line) indicates main differentiation trajectory, lineage 2 (dashed green) indicates basal cell renewal pathway. G) Pseudotime distribution of basal, suprabasal, atypical and ciliated cell population densities across the major lineage 1 identified in (F), showing the differences in cell type proportions between the three treatments (Kolmogorov-Smirnov test, Control versus NTCU 10µM, *p*<0.001). H) Heatmap showing scaled expression of the top 75 marker genes of the atypical (ABC) cluster. Gene-level significance was assessed using Seurat FindAllMarkers: a two-sided Wilcoxon rank-sum test with Bonferroni correction, and genes are ranked based on average log_2_ fold change. Expression values represent z-scored log-normalised data and capped at ±2.5 to improve contrast. A random sample of 2000 cells are plotted in each column, cell type is indicated above. I) Violin plots showing the distribution of *VIM* log-normalised counts in cells from each annotated cluster, coloured by treatment condition (Control, 3µM, 10µM). Differential expression analysis was performed using Seurat, *VIM* was the second most significant marker gene of the atypical (ABC) cluster (adj. *p*<0.001, Wilcoxon rank-sum, Bonferroni corrected). *VIM* was also upregulated in basal cells (basal, suprabasal, cycling basal) in NTCU 10µM versus Control (adj. *p*<0.001, Wilcoxon rank-sum, Bonferroni corrected). J) IF staining of Control and NTCU-treated ALI cultures for VIM (green), with nuclei counterstained with Hoechst (blue). Scale bar, 150µm. Representative images from three biological replicates. K) Enrichment plot showing enrichment of the Hallmark EMT geneset in basal cells (including basal, cycling basal and suprabasal) treated with NTCU (10µM) versus Control (Benjamini-Hochberg adj. *p*=0.0019). Gene set enrichment analysis (GSEA) performed on ranked log_2_FC values, using the fgsea algorithm. L) Heatmap showing scaled expression of top 20 genes from the Hallmark EMT geneset, for basal cells (including basal, cycling basal and suprabasal) treated with Control, 3µM NTCU and 10µM NTCU. Genes selected based on ranked log_2_FC values in 10µM NTCU versus Control. Values scaled as in (H). A random sample of 2000 cells are plotted, treatment (Control and NTCU dosage) is indicated above. M) Violin plots showing the distribution of *COL1A1 (adj. p<0.001), FN1 (adj. p<0.001), VCAN (adj. p<0.001) and TWIST (adj. p<0.001)* log-normalised counts in cells in basal cells (basal cells only), split by treatment condition (Control, 3µM, 10µM). Two-sided Wilcoxon rank-sum test, Bonferroni correction for multiple testing, comparing expression in basal cells treated with Control versus 10µM NTCU.

To examine the impact of the carcinogen, we next visualised the single cell data split by treatment and patient sample (**Figure 2B**). There was a dose-dependent expansion of the atypical cell population in the NTCU-treated samples compared to controls, suggesting the emergence of these cells is a direct response to NTCU exposure. We confirmed this cell population was treatment-specific by analysing the control and NTCU-treated groups separately showing that the atypical population was unique to the treated samples (**Figure 2C**).

The novel population accounted for 12.8% of the epithelial cells in the 10μM NTCU treatment. This expansion (*p*=0.0082) was accompanied by a corresponding decline in the *KRT5^+^* basal cell population (*p*=0.0022; **Figure 2D, Supp. Figure 1B**), suggesting a basal origin for the atypical cells, and consistent with prior work on the cell-of-origin of murine LUSC^9^.

To explore this, we applied two trajectory approaches: Monocle3^18^ and Slingshot^19^. Both tools order cells along a pseudotime axis by progressive changes in gene expression, with Slingshot additionally resolving branching lineages representing distinct cell fate paths.

Monocle3 defined a biologically plausible trajectory consistent with the atypical cells being derived from basal cells (**Figure 2E**). Slingshot defined two lineages: (1) from basal through atypical to ciliated cells, and (2) a self-renewal pathway from basal to proliferating basal and back again (**Figure 2F**). Along Lineage 1, pseudotime density revealed a dose-dependent shift with NTCU exposure (*p*<0.001): control cells clustered at early (basal) pseudotime, whereas treated cells accumulated at later values, reflecting transition towards the atypical cell state **(Figure 2G and Supp. Figure 1C**).

To further validate this without imposing any prior assumptions about cell of origin, we applied RNA velocity analysis using scVelo^20^, which infers the direction of cell state transitions from transcriptional kinetics. This orthogonal approach was again consistent with the atypical cells representing a transitional population derived from basal cells (**Supp.Figure 1D**). Taken together the evidence supports the notion that NTCU-derived atypical cells in this model are basal cell derived - “atypical basal cells” (ABCs), consistent with what is hypothesised about human LUSC and what is known from mouse models^9^.

Next, we examined the transcriptional profile of the ABCs. Using Seurat’s *FindAllMarkers* protocol, we identified 372 genes (**Supp. Table 3**) specifically upregulated in the ABC population (**Figure 2H**). The ABCs were defined by upregulation of genes spanning several cancer-associated programmes (Figure 2H, Supp. Table 3). These included genes associated with epithelial-mesenchymal transition (EMT), plasticity and invasion (*VIM*, *MACC1*, *SOX9*, *SERPINB2*, *ANXA1*, *ANXA3*); an altered keratin profile (Keratin-7 (*KRT7*), *KRT8*, *KRT23*, *KRT17*); and cell-cycle checkpoint and senescence regulators, several of them p53-network genes (*CDKN1A* [p21], *CDKN2A* [p16], *MDM2*, *SFN* [14-3-3σ]).

Of these marker genes, *VIM* had one of the largest log fold changes compared to all other cell types (adj. *p*<0.001; **Figure 2I**, **Supp. Figure 1E**). *VIM* is generally associated with fibroblast/stromal cells rather than epithelial cells and is a canonical marker of EMT, a critical early event in epithelial carcinogenesis^21^. We used immunofluorescence (IF) to confirm *VIM* upregulation in the carcinogen-exposed cells (**Figure 2J**).

We also noted that basal cells upregulated *VIM* on treatment with NTCU (*p*<0.001), suggesting that carcinogen exposure upregulates EMT-related genes in the transition towards the ABC state. This was confirmed by a Gene Set Enrichment Analysis (GSEA) comparing control and NTCU-treated basal cells (basal, cycling basal and suprabasal, adj. *p*=0.0019; **Figure 2K, Supp. Figure 1F**) and included a dose-dependent upregulation in basal cells of key EMT genes such as *TWIST* (adj. *p*<0.001), a canonical EMT transcription factor and COLA1 (adj. *p*<0.001), *FN1* (adj. *p*<0.001) and *VCAN* (adj. *p*<0.001; **Figure 2L-M**). There was therefore a marked change in cell state in response to carcinogen, consistent with early precancerous change.

### ABCs are characterised by key markers of metaplasia and by activation of the DNA damage response

In the paradigm of lung squamous carcinogenesis, there is a stepwise progression from normal epithelia through hyperplasia/metaplasia and then progressive dysplastic change. The change from normal pseudostratified epithelium to metaplasia is a critical early event; metaplasia has been demonstrated in examples of early epithelial carcinogenesis associated with lineage plasticity^22,23,24^. This plasticity is often associated with *SOX9* expression and/or alterations in keratin expression patterns^22,24,25^.

*SOX9* was upregulated in the ABC population after exposure to NTCU (adj. *p*<0.001; **Figure 3A, Supp. Figure 2A, Supp. Table 3**), and increased expression confirmed by immunohistochemistry (IHC) **(Figure 3B)**. This was associated with a downregulation in *SOX2* expression. *SOX2* is implicated as a lineage driver oncogene later in LUSC pathogenesis^26,27^ so this may be counterintuitive but, in fact, *SOX2*-*SOX9* reciprocal regulation is well described. Further, an early reduction in *SOX2* prior to later deregulated overexpression was previously hypothesised in LUSC precancerous progression^28^. The emergence of ABCs provides evidence in support of that hypothesis.

**Figure 3.**
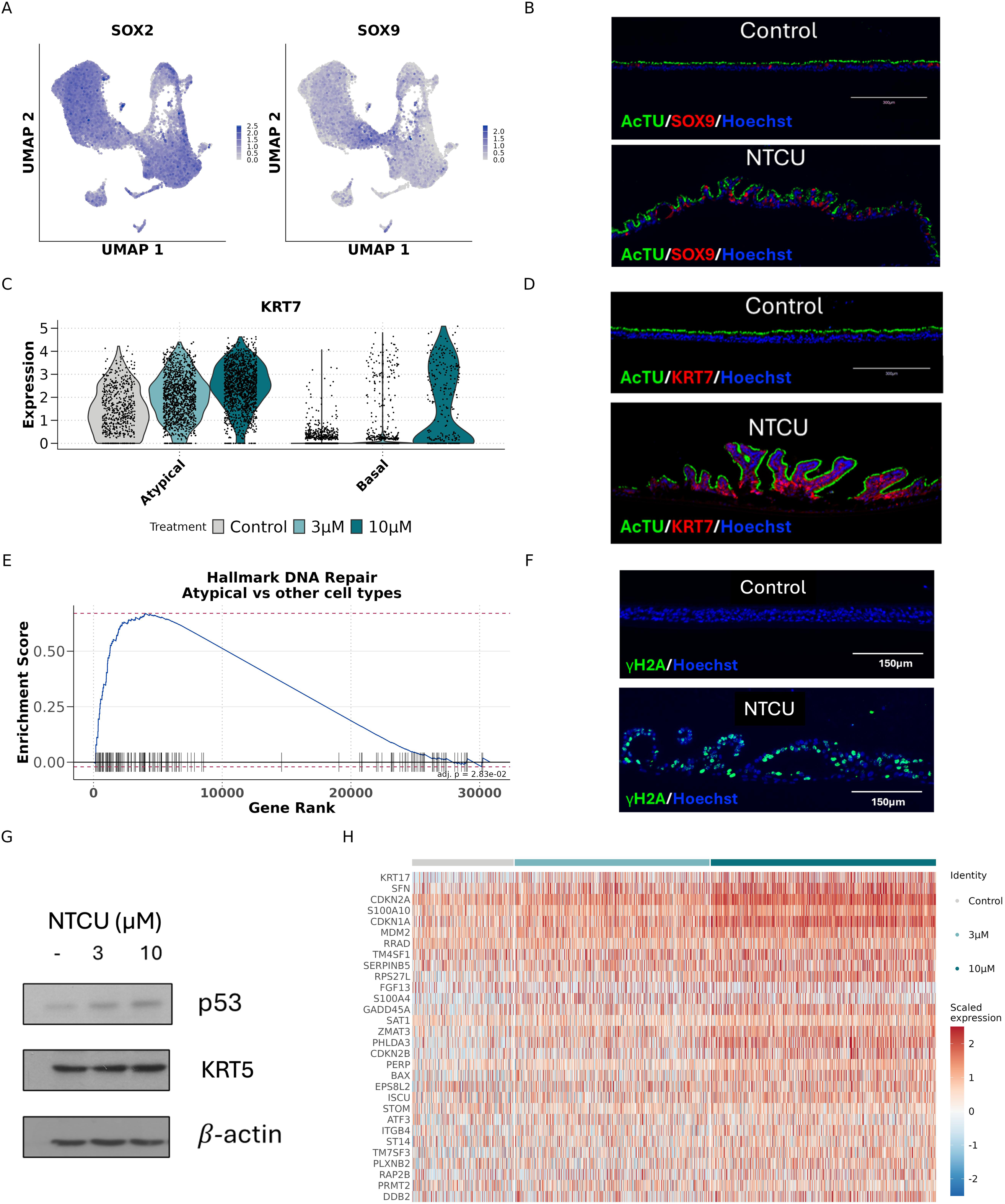
Expression of markers of metaplasia and the DNA damage response in ABCs. (A) UMAPs depicting expression of *SOX2* and *SOX9*. (B) IF staining of paraffin-embedded sections of Control and NTCU-treated ALI cultures, for SOX9 (red), acetylated tubulin (green) and Hoechst nuclei counterstain (blue). Image is representative of three replicates. Scale bar, 300µm. (C) Violin plot showing the distribution of *KRT7* log-normalised counts in cells from the basal and atypical (ABC) clusters, coloured by treatment condition (Control, 3µM NTCU, 10µM NTCU). Increase in KRT7 expression in ABCs in NTCU versus Control, adj. *p*<0.001, Wilcoxon rank-sum, Bonferroni corrected. (D) IF staining for KRT7 (red), acetylated tubulin (green) and Hoechst nuclei counterstain (blue) in Control (top) and NTCU 30µM (bottom). Scale bar, 300µm. Image is representative of three replicates. (E) Gene set enrichment profile for the Hallmark DNA Repair geneset, comparing gene expression between atypical (ABC) cells and all other cell types (adj. *p*=0.028). GSEA performed on ranked log_2_FC values, using the fgsea algorithm. Running enrichment score is shown across the ordered gene, vertical marks indicate the positions of DNA Repair pathway genes. (F) IF staining of γH2AX (green) and Hoechst nuclei counterstain (blue) in Control (left) and NTCU 10µM (right) treated ALI cultures. Scale bar, 150µm. Image is representative of five replicates. (G) Western blot of p53 and KRT5 expression in human bronchial epithelial cell (HBECs) cultured at ALI following treatment with Control, 3µM or 10µM NTCU. Western blot representative of two experiments. (H) Heatmap showing scaled expression data for the top 30 upregulated genes from the Hallmark p53 pathway geneset in atypical (ABC) cells (ranked by average log_2_FC versus all other cell types), split by treatment. A random sample of 2000 atypical (ABC) cells are plotted, treatment indicated above.

*KRT7* is a target of SOX9 and a key marker of SOX9-associated metaplasia^24^. Together with *VIM*, *KRT7* was elevated in the ABC population (*p*<0.001; **Figure 2H; Supp. FIgure 2B, Supp. Table 3)**. In response to NTCU treatment there was a dose-dependent increase in transcription of *KRT7* in the ABC population (*p*<0.001), along with the basal population, confirmed using **IF** (**Figure 3C & D**). *KRT17* expression has also been linked to metaplastic change^29,30^ and was significantly enriched in the ABC population and increased in response to NTCU (both adj. *p*<0.001; **Supp. Table 3**, **Supp. Figure 2B-D**). The combination of EMT and these markers of metaplasia indicates the ABCs have an abnormal plastic cell state consistent with the earliest stages of squamous carcinogenesis.

When comparing the ABCs with all other cell types the most significantly upregulated pathway was DNA repair (adj. *p*=0.028; **Figure 3E, Supp. Figure 2E)**. This was associated with marked induction of γH2AX, a canonical marker of the cellular response to DNA double-strand DNA breaks, and stabilisation of p53 (**Figure 3F & G, Supp. Figure 2F & G**). We saw an upregulation of p53 response genes in ABCs and observed a dose-dependent upregulation in response to NTCU exposure, (**Figure 3H**), including key mediators such as *BAX*, *ZMAT3*, *SFN* and *MDM2*. These observations are consistent with NTCU acting as an alkylating agent that directly damages DNA and is typical of a carcinogen-induced model of precancer.

Changes in cell state have been associated with lung pathology, notably human idiopathic pulmonary fibrosis (IPF), in which a specific, consistent cell population with an abnormal transcriptional profile has been described, with similar populations reported in murine models of IPF ^31–33^. We compared the ABC population with the IPF-associated cell states (**Supplementary Figure 3A-C**). There were similarities in terms of altered keratin expression - upregulation of KRT17, KRT7, and KRT8; and the upregulation of markers associated with EMT programmes and the DNA damage response. This is consistent with shared lung epithelial responses to external injury. However, there were also marked differences, likely reflecting the origin of the aberrant basaloid cells from alveolar type 2 (AT2) cells and the distinct pathogenic programmes associated with a fibrotic process.

We next compared the ABCs to scRNASeq datasets from tracheal brushing from smokers and to murine hillock cells (**Figure 3D & E**) ^9,34^. Hillock cells have recently been described in humans and implicated in metaplastic change in the context of vitamin A deficiency ^34^. We also noticed that SERPINB2, was a key marker of murine hillock cells and a marker of the ABCs. However, in general there was limited shared transcriptional identity with the ABCs.

Taken together, this analysis is consistent with the basal cell-derived ABC cell state being distinct from that of the IPF-associated aberrant basaloid cell and distinct from either normal hillock cells or cigarette smoke-exposed normal tracheal cells.

### Carcinogen-induced phenotypic changes are not associated with typical mutational driver events or dominant clonal expansion

We next addressed whether the NTCU-induced phenotype was associated with the typical mutational events known to be prevalent in the early stages of squamous carcinogenesis. We also examined whether there was evidence of broader genome disruption detected by whole genome sequencing.

We compared DNA extracted from differentiated airway cultures exposed to vehicle control or NTCU for six weeks. We performed high-coverage sequencing of mutational hotspots in 46 key driver genes implicated in solid organ malignancies including LUSC (**Figure 4A**). The targeted genes are listed in **Supp. Table 4,** and are part of a typical clinical targeted-sequencing panel.

**Figure 4.**
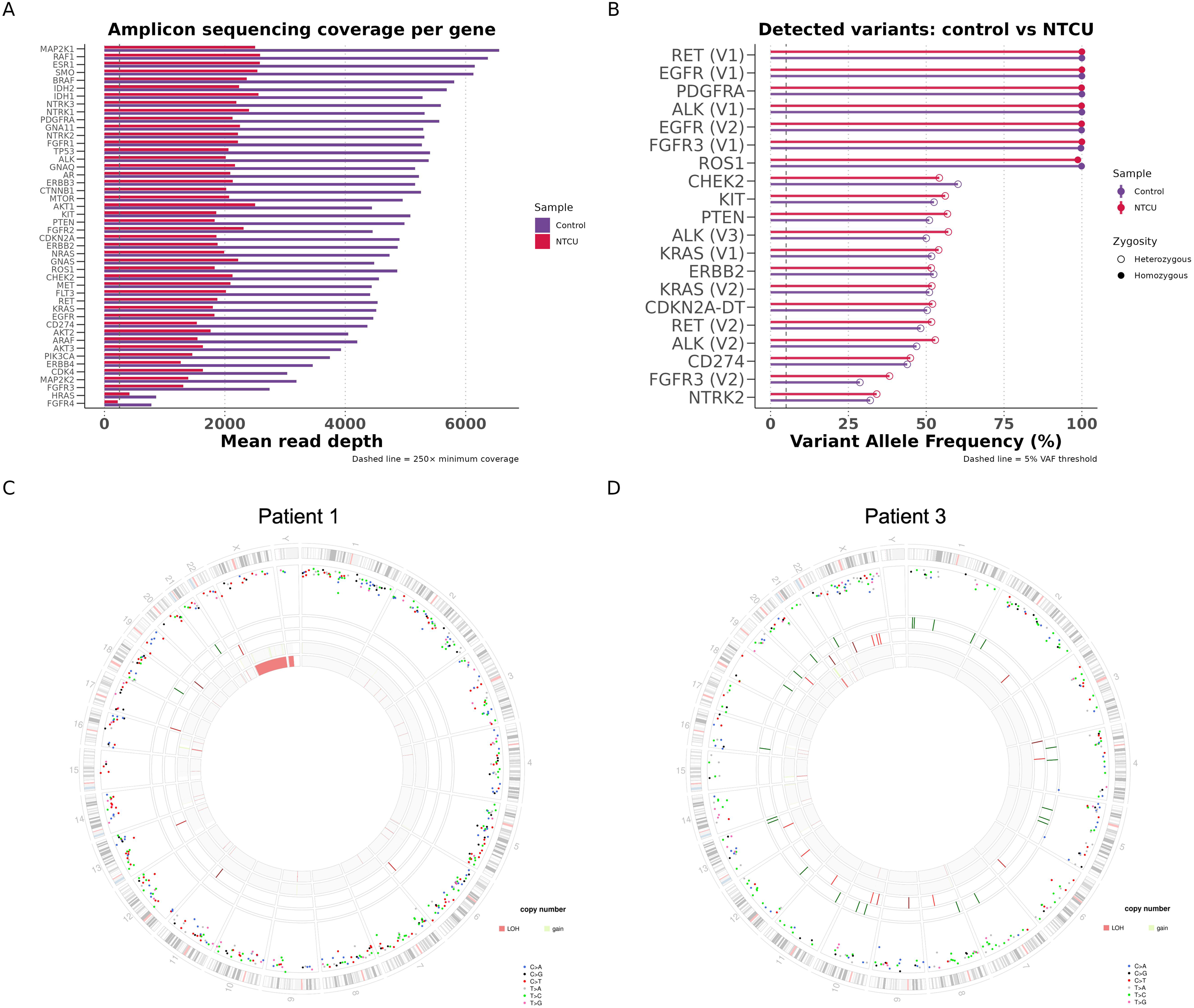
Genomic characterisation of the NTCU model. (A) Mean read depth across all amplicons per gene in the 46-gene cancer hotspot panel. Values are shown for both Control and 10µM NTCU-treated cultures from a single patient sample (Patient sample 3). Dashed line indicates 250× target coverage. (B) Variant Allele Frequencies (VAFs) of all variants detected above the threshold, in either treatment. Filled and open circles denote homozygous and heterozygous calls, respectively. Dashed line indicates the 5% minimum detection threshold. y axis labels indicate gene followed by variant number, in cases where multiple variants were detected in the same gene. *CDKN2A-DT* refers to the divergent transcript lncRNA at the *CDKN2A* locus. (C)(D) Circos plot from whole-genome sequencing comparing patient samples (1 and 3) treated with 10µM NTCU vs Control. Each circos plot is organised from the outermost ring inwards as follows: chromosomal ideogram; Single Bases Substitutions (SBSs) displayed as a rainfall plot (log_10_ intermutation distance on the radial axis; C>A, blue; C>G, black; C>T, red; T>A, grey; T>C, green; T>G, pink); small insertions (green ticks) and deletions (red ticks); major copy number allele (green, gain); minor copy number allele (red, loss).

Mean amplicon coverage exceeded 250X in 45 of 46 genes in both control and NTCU-exposed cultures. No somatic events were detected at this resolution (**Figure 4B)**, therefore excluding dominant clonal (or significant subclonal) mutations in genes implicated in LUSC pathogenesis including *CDKN2A*, *TP53* and *PTEN*, as driver events in our model.

Whole genome sequencing (WGS) was also used to screen for large-scale genomic events (**Figure 4C,D**). Despite the evidence of double strand breaks (DSBs) reflected by the engagement of the DNA damage response (γH2AX, **Figure 3E-H**), the genomes of the NTCU-exposed epithelioids were not grossly abnormal. There was a modest increase in single base substitutions (SBSs) and insertion/deletions (indels) in both patient samples after exposure to NTCU (**Supp. Figure 4A-D**). Although we acknowledge the limitations of WGS at this coverage (20-30x) in terms of robust discovery of rare variants, the data are consistent with there being no evidence of a dominant clonal expansion associated with the NTCU phenotype. Given there was not a dominant clonal population, mutational signature analysis was not informative.

Taken together, the carcinogen drives a basal cell phenotype consistent with the earliest stages of squamous carcinogenesis, but without evidence of a clonal or dominant subclonal expansion. We next compared the model with available clinical samples.

### The NTCU-exposed ALI model recapitulates key expression patterns described in early lung squamous carcinogenesis

To assess the clinical relevance of the NTCU exposure model, we interrogated an independent, publicly-available gene expression dataset of human bronchial biopsies spanning the full histological spectrum of lung squamous carcinogenesis, generated by Mascaux et al^35^. This comprised 122 clinical specimens from 77 patients. Specifically, the cohort included basal cell hyperplasia, squamous metaplasia and low-grade dysplasia - the putative earliest stages of squamous carcinogenesis.

We created an “ABC signature” from the top 20 altered ABC marker genes (defined in Figure 2, ranked by average log_2_ fold change (log_2_FC)) in ABCs from NTCU-exposed cultures and mapped that signature onto the independent series. The results demonstrate an increase in the average expression of the ABC signature with histological progression to moderate dysplasia (Spearman coefficient=0.4, *p*<0.001; **Figure 5A)**.

**Figure 5.**
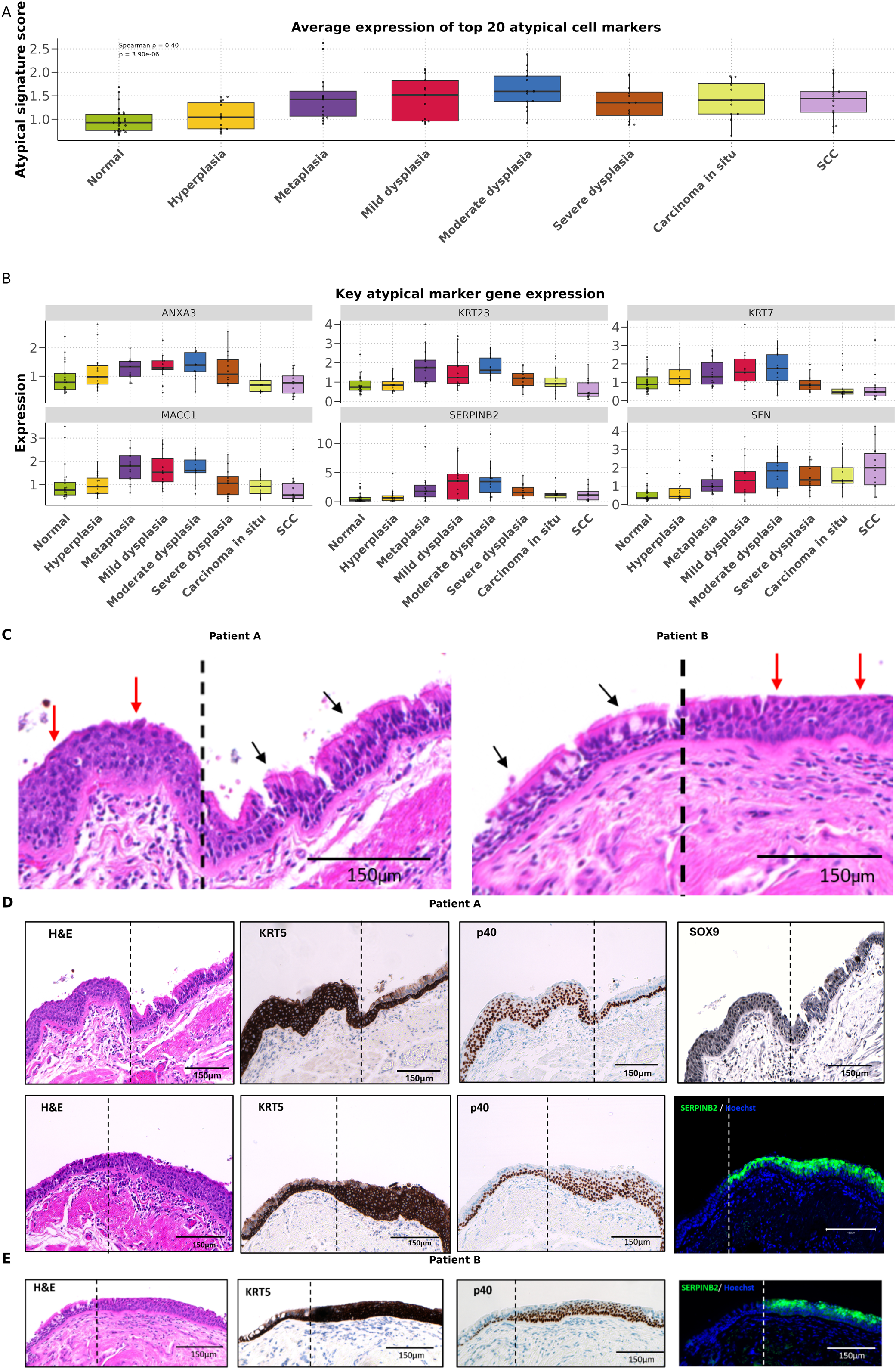
ABC marker genes are upregulated in early bronchial preneoplasia. (A) Atypical (ABC) signature score (mean expression of the top 20 ABC marker genes) across eight histological stages in the Mascaux et al. bronchial biopsy microarray dataset. Spearman correlation with stage is shown, *p*<0.001. Each point represents one biopsy; boxes show median ± interquartile range (IQR). Sample sizes as follows: Normal (27), Hyperplasia (15), Metaplasia (15), Mild dysplasia (13), Moderate dysplasia (13), Severe dysplasia (12), Carcinoma in situ (13), Squamous cell carcinoma (SCC) (14). (B) Expression of six key atypical (ABC) marker genes (*KRT7, MACC1, ANXA3, SERPINB2, SFN, KRT23*) across the eight histological stages. Each point represents one biopsy; boxes show median ± IQR. (C) Histological specimens from two patients (A and B) in which the junction between normal pseudostratified epithelium (black arrows) and metaplastic/low-grade dysplasia (red arrows) was identified. Scale bar, 150µm (D) Sections through the proximal and distal junctions between normal pseudostratified epithelium and a low-grade dysplastic lesion from Patient A. IHC (KRT5, p40, SOX9) and IF (SERPINB2, green). Scale bar, 150µm (E) H&E of the pseudostratified-low-grade dysplasia junction from Patient B. IHC (KRT5, p40) and IF (SERPINB2, green). Scale bar, 150µm

We next selected individual markers that contribute to the signature to illustrate the link with histological progression (**Figure 5B)**. In each case there was an increase in expression at the earliest stages of the disease process. One of the markers - *KRT7* - we have already interrogated as a marker of cell state change (**Figure 3**). The other markers include proteins linked with the DNA damage response^36^ or implicated in cancer progression^37,38^. As a negative control we compared the median ABC signature score to 20 randomly sampled gene sets of equivalent size (n=20) across histological stages, to show that this pattern of increase to metaplasia is specific to the ABC marker genes (**Supp. Figure 5A**).

We then identified three patients (A, B and C) who have undergone surgery for non-small cell lung cancer and for whom we had captured the junction between normal bronchial epithelium and metaplasia within the resection specimen. Consistent with the carcinogen model and the independent precancer transcriptome dataset, there was evidence of an expansion of KRT5^+^ / p40^+^ cells that were also positive for SOX9 (Patient sample A) and SERPINB2 (Patient samples A, B and C) (**Figure 5C-E, Supp. Figure 5B)**. p40 recognises a truncated form of p63 (ΔNp63) and is used more commonly in clinical practice.

### Capivasertib treatment prevents and reverses carcinogen-induced phenotype

Capivasertib, a multi-AGC kinase/pan-AKT inhibitor used clinically in combination with fulvestrant for metastatic breast cancer and under consideration for combination treatment of metastatic prostate cancer, was previously shown to have efficacy in preventing high-grade bronchial dysplasia in a SOX2-dependent organotypic model^39^. We therefore tested the impact of capivasertib in the NTCU model, mimicking the potential for chemoprevention early in LUSC carcinogenesis.

The initial experimental set-up is shown in **Figure 6A**. Capivasertib was added to differentiated airway epithelioids at the same time as the carcinogen. Treatment with carcinogen was continuous but for capivasertib was for four days on and three days off to replicate the clinical dosing regimen. We demonstrated the expected pattern of target engagement (**Supp. Figure 6A**). Remarkably, for each of three patient samples, capivasertib prevented the emergence of the NTCU phenotype with preservation of pseudostratified airway histology (**Figure 6B, C**). The concentrations used (range 1.5-3µM) are achievable with the standard oral dose used in the clinic^40,41^.

**Figure 6.**
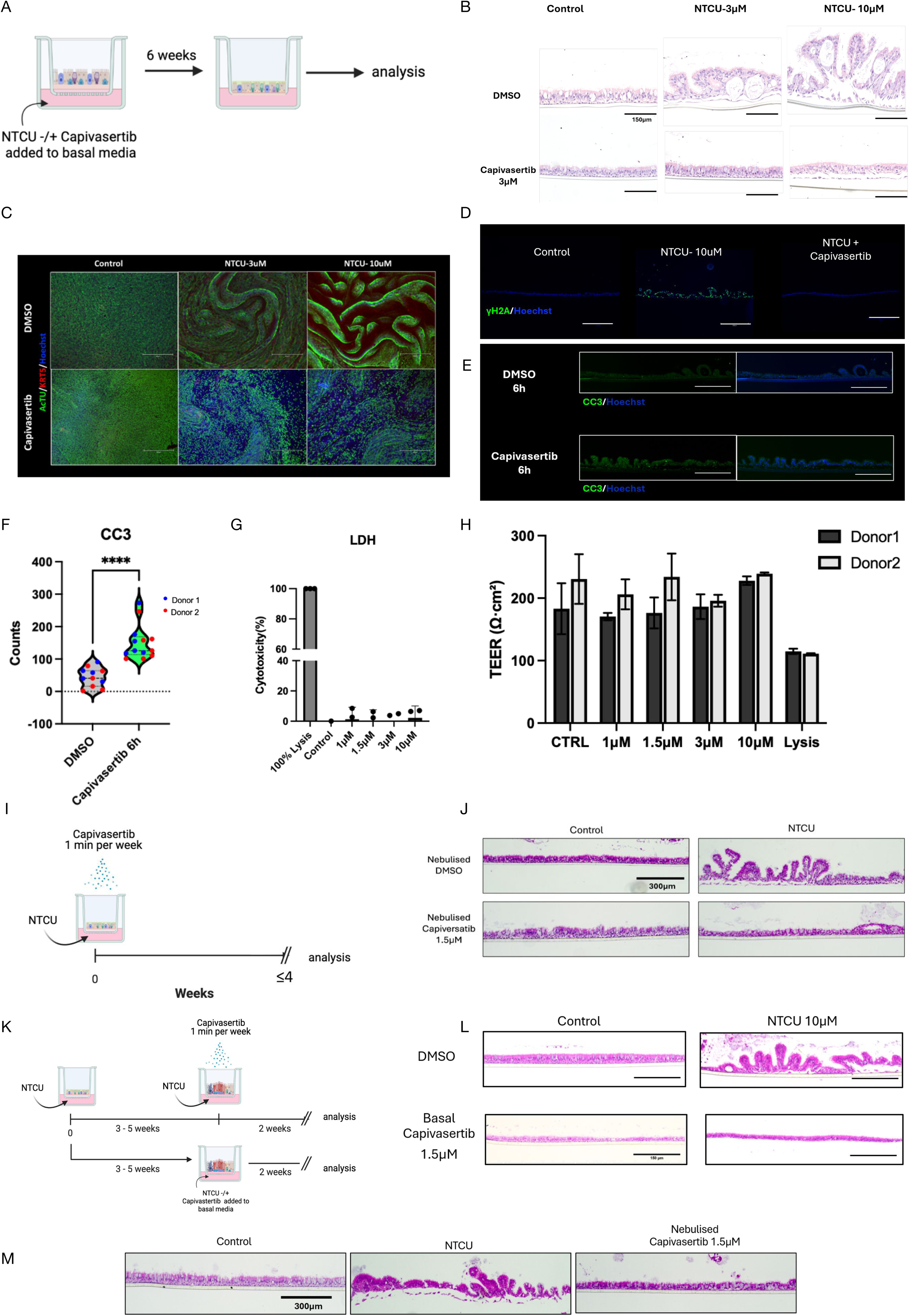
Capivasertib prevents and reverses NTCU-induced premalignant airway lesions in human ALI cultures. (A) Schematic overview of the experimental design. Primary human airway epithelial cells cultured at ALI were exposed to NTCU (3 or 10μM) added to the basal medium in the presence or absence of the AKT inhibitor capivasertib (3μM). Cultures were maintained for six weeks before analysis. (B) Representative H&E-stained sections of ALI cultures following treatment with vehicle control (dimethyl sulfoxide, DMSO), NTCU (3μM), or NTCU (10μM), with (bottom) or without (top) capivasertib (3μM). NTCU induced epithelial hyperplasia and dysplastic-like morphological changes, which were attenuated by capivasertib treatment. Scale bars, 150μm. Representative images from two replicate experiments on cells from two patient samples. (C) Representative IF images showing epithelial differentiation and squamous marker expression in ALI cultures treated with vehicle control, NTCU (3μM), or NTCU (10μM), with (bottom) or without (top) capivasertib (10μM). Sections were stained for acetylated tubulin (green), KRT5 (red), and Hoechst (blue). Scale bars, 300µm. Representative images from two replicate experiments on cells from two patient samples. (D) Representative IF staining for γH2AX (green) and Hoechst (blue) demonstrating DNA damage in Control, NTCU-treated (10μM), and NTCU plus capivasertib-treated (10μM) cultures. Co-treatment with capivasertib prevented the accumulation of γH2AX^+^ cells. Scale bars, 150µm. Representative images from three replicate experiments on cells from two patient samples. (E) Representative IF images of cleaved caspase-3 (CC3; green) and Hoechst (blue) staining of NTCU-treated ALI cultures following six hour treatment with DMSO (top) or capivasertib (10μM). Increased CC3 staining indicates induction of apoptosis in capivasertib-treated cultures. Scale bars, 150µm. Representative images from two patient samples. (F) Violin plot of CC3^+^ counts following 6 hour treatment with DMSO or capivasertib; individual points represent the quantification from a single image, coloured by donor (blue, Donor 1; red, Donor 2). Dashed line, median; dotted lines, quartiles. CC3-positive cells following six hour treatment with capivasertib or DMSO control were quantified using Fiji by measuring the green fluorescence signal in each image. **** = *p*<0.0001 (Student’s *t*-test). (G) Lactate dehydrogenase (LDH) release assay measuring cytotoxicity (%) in differentiated ALI cultures (in the absence of NTCU) treated with basal capivasertib at 1, 1.5, 3 and 10 µM, alongside a vehicle control and a 100% lysis positive control. Bars show mean ± s.d. and each point represents one patient sample (n=2 patient samples). Note the discontinuous y-axis. (H) Transepithelial electrical resistance (TEER; Ωcm²) of differentiated ALI cultures treated with basal capivasertib in the absence of NTCU (1, 1.5, 3 and 10 µM), an untreated control and a positive lysis control. Bars, mean ± s.d.; n = 2 patient samples. (I) Schematic of the experiment modelling the efficacy of nebulised (“inhaled”) capivasertib in preventing the NTCU phenotype. Healthy differentiated ALI cultures were exposed simultaneously to continuous basal NTCU (10µM) along with nebulised 1.5-3µM Capivasertib for one min each week for three or four weeks. (J) Representative H&E-stained sections of ALI cultures treated with Control and 10μM NTCU, and with nebulised DMSO or nebulised capivasertib. Capiversatib treatment prevented the NTCU-induced phenotype. Scale bar, 300μm. Representative histology from experiments on three patient samples (1.5µM, n=2; 3µM, n=2). See Supplementary Figures for representative histology from experiment using 3µM. (K) Schematic of experiment aimed at testing the efficacy of capivasertib in reversing the NTCU phenotype with capivasertib delivered basally or nebulised. NTCU exposure was performed for up to five weeks prior to either basal DMSO or capiversatib treatment (four days on, three days off) or nebulised DMSO or capiversatib treatment (one min per week) for an additional two weeks before analysis. (L) Representative histological images from intervention experiment depicted in (K), showing airway epithelial morphology following NTCU exposure with or without basal capivasertib treatment. Scale bar, 300μm. (M) Representative histological images from intervention experiment depicted in (K), showing airway epithelial morphology following NTCU exposure with or without nebulised capivasertib treatment. Scale bar, 300μm.

We showed above that NTCU induces the DNA damage response in this culture system (**Figure 3**). We therefore interrogated the impact of capivasertib and found that it prevented the accumulation of cells positive for γH2AX (**Figure 6D)**, a key goal of a successful chemoprevention regime. Capivasertib has previously been shown to induce apoptosis in a minority of cells vulnerable to the drug^42^. We generated cultures with the NTCU phenotype and measured the impact of capivasertib using cleaved caspase-3 (CC3) as a marker of apoptosis (**Figure 6E)**. We demonstrated a significant increase (*p*<0.001) in apoptosis six hours after treatment commenced, illustrating a vulnerability of NTCU-exposed cultures to this targeted therapeutic (**Figure 6F**).

Potential toxicity is a key consideration for any proposed chemoprevention. The effective *in vitro* dose is achievable *in vivo*. Further, thousands of patients have now been treated with capivasertib without overt pulmonary toxicity, which is reassuring^43,44^. However, to directly assess this we exposed differentiated ALI epithelioids to increasing concentration of capivasertib and used standard airway toxicity assays to measure its impact. There was no reduction in Transepithelial Electrical Resistance (TEER) or increase in lactate dehydrogenase (LDH) release as a result of capivasertib treatment **(Figure 6G & H)**.

Finally, we appreciate that systemic (oral) capivasertib is not an attractive clinical approach to the chemoprevention of lung cancer due to the systemic side-effect profile (notably rash, diarrhoea and hyperglycaemia) ^45,46^ lower therapeutic index of orally compared to inhaled delivered drugs for the treatment of lung diseases ^47^. We therefore tested the potential to nebulise (“inhale”) the drug (**Figure 6I**). We titrated the dose so that the mass of capivasertib delivered (0.45 nmol = 193 ng) was similar (2-3-fold lower) to the therapeutically effective basal (“systemic”) treatment with 1.5-3 µM in 500 µl of basal medium (0.75 - 1.5 nmol; 321 - 642 ng). This was achieved by administering 0.25 μl of nebulised 1.8 mM, although nebulisation took place for only 1 minute/week rather than the systemic (basal) regime of 4 days-on/3 days-off ^48,49^. Similar to previous studies with small molecule targeted inhibitors ^50^, we found that nebulised capivasertib was equally effective at preventing the NTCU-phenotype even at the lower dose, but with orders of magnitude less of total dose under clinical conditions as discussed below (**Figure 6J, Supp. Figure 6B**).

Further, we wanted to test the potential of capivasertib to reverse the NTCU-phenotype once established (**Figure 6K)**. Remarkably, we found that both basal (systemic) (**Figure 6L**) and nebulised (inhaled) (**Figure 6M**) capivasertib were effective suggesting a potential role for nebulised small molecules in both primary and secondary chemoprevention of lung cancer.

The potential advantage of the inhaled compared to oral delivery is readily achieving effective local therapeutic concentrations despite substantially lower doses. In the case of salbutamol, experimental data in volunteers demonstrated a 2400-fold lower airway concentration after oral dosing compared to dose-normalised inhalation ^51^. We do not have similar data for capivasertib but did perform *in silico* modelling of the likely pulmonary availability of inhaled capivasertib using a custom-built software which has been validated with clinical data ^52^. This predicted, using an inhaled bd regimen (twice daily) of 100µg as an example, that a concentration of >3µM could be achieved in the epithelial lining fluid - equivalent to approximately 10^5^ larger than that achieved by dose-normalised oral dose. At this dose the systemic concentration is predicted to be <3nM, a concentration not expected to result in systemic side effects.

## Discussion

In this work we expose long-term differentiated multicellular pseudostratified airway epithelia to a carcinogen. The carcinogen directly impacts the basal cell population, altering basal cell dynamics and leading to a novel basal cell-derived cell state that we term atypical basal cells (ABCs). These cells exhibit hallmarks of cell plasticity that are also hallmarks of cancer and share core transcriptional signatures with the earliest stages of human LUSC. We then show that we can prevent and reverse the carcinogen-associated histological phenotype by repurposing a drug already in the clinic. Further, the data suggests that the early precancer phenotype is not the result of a dominant clonal or subclonal expansion, consistent with it not being mediated by a specific mutational event.

LUSC has lagged behind lung adenocarcinoma in terms of therapeutic innovation and clinical outcomes. This, in part, reflects the absence of rational and appropriate models of the disease that allow the pathobiology to be probed and novel therapeutic approaches to be tested. There is rightly a renewed focus on rational non-animal models of disease, particularly those using primary human cells, as a result of recent initiatives from the Federal Drug Administration and the European Medicines Agency^53^. Models such as those presented here are therefore of increasing importance for translation to the clinic. Further, there is an increasing emphasis on earlier intervention and prevention to reduce cancer morbidity and mortality.

Exposure to NTCU disrupts basal cell homeostasis and drives the emergence of ABCs with a unique transcriptional signature. Importantly, this signature was reproduced in an independent transcriptional dataset derived from early human precancer LUSC biopsies. The model therefore is representative of precancerous LUSC and this adds confidence to our interpretation of the impact of capivasertib.

The altered transcriptome and cell trajectory analyses suggest an evolving cell state under carcinogen pressure. In particular, the emergence of a *SOX9*^+^ population is consistent with what is known about its role as a master controller of cell state, including specifically in metaplastic change^25,54^. In LUSC a metaplastic change is a prerequisite for the progression from a pseudostratified epithelium to squamous dysplasia and then invasive LUSC. Metaplasia is commonly associated with EMT or epithelial-mesenchymal plasticity (EMP) programmes and these were activated in ABCs and basal cells in response to NTCU. The plasticity implied by the EMT/EMP continuum is consistent with the clinical potential of preinvasive lesions to both progress and regress^55^.

Interestingly, the ABCs share some key markers with aberrant basaloid or KRT5-/KRT17+ cells recently reported in IPF^3132^ in which *SOX9* and an EMT/EMP programme are also implicated. Further shared markers include *CDKN1A*, *CDKN2A*, *MDM2*, and *GDF15*, a gene set associated with DNA damage and cancer. However, there are also major differences including the repertoire of fibrosis-associated genes expressed in the IPF-associated cell state but not ABCs (including *AVB6* (*ITGB6*), *MMP7*, *EPHB2*), as well as well as the putative cell of origin in IPF being the AT2 cell ^56^.

It is of note that the ABC transcriptional signature includes *SERPINB2*. Interestingly, Mascaux and colleagues reported two biphasic “gene modules” in human LUSC precancer specimens, including one characterised by *SERPINB2* and other SERPIN family members^35^. SerpinB2 was recently identified as a marker of airway hillock cells in the mouse along with Krt13/KRT13 in both the mouse and human ^34^. Hillock cells are an important population of cells that are injury resistant reservoirs of stem cells with the capacity to repopulate damaged airway pseudostratified epithelium and that can give rise to squamous metaplasia in the context of vitamin A deficiency in the mouse^56^. The cell trajectory analysis and clonal expansion in the mouse NTCU-precancer/LUSC model suggested the Krt13^+^ pathogenic clones in their work were derived from a Krt5^+^ pool rather than necessarily the hillock subpopulation^9^. The ABCs in our human NTCU-exposure model are derived from the *KRT5^+^* population but do not express *KRT13*. These differences may reflect the different species, as well as the pre-clonal events being modelled in this work.

The impact of a carcinogen in driving an abnormal phenotype, associated with alterations in basal cell dynamics and cancer hallmarks events (DNA damage, EMT, plasticity, metaplasia) is notable as it suggests that NTCU changes cell state in the absence of a clonal expansion. There is accumulating evidence that carcinogens can act by tumour promotion rather than mutation, potentially on the background of previously acquired mutations^57^. For example, recent work on murine cutaneous squamous cell carcinoma described a cell state evolution in mouse skin in response to mutagen exposure followed by protracted tumour promotion. Our data suggest that comparatively short-term exposure to a carcinogen in differentiated human cells confers cell state changes typical of precancer, but with no evidence of prior or emergent clonal mutations in driver genes.

Taken together, our data is consistent with the notion that airway cell lineages, in this case basal cells, respond to insult in a limited number of ways and that the plastic cell state that emerges in response to NTCU expands a population of cells that may be vulnerable to further insult or mutational events.

This work has some limitations. Despite best efforts we were unable to establish the culture conditions to propagate the ABCs from these complex epithelial models, similar to the challenges described with the aberrant basaloid cells identified in idiopathic pulmonary fibrosis^31^. This also reflects the challenges in propagating patient-derived organoids in LUSC. An expanded pool of ABCs would be desirable for functional characterisation and to probe mutational signatures and events in more detail. We also acknowledge that, although NTCU generates a compelling model that is experimentally tractable, the duration of carcinogen exposure in the human airway and the natural history of developing precancer are much more protracted. We acknowledge the significant challenges associated with proving the absence of a mutational event in a polyclonal population. It is possible that there are many low frequency subclonal events below the reported sensitivity of the panel used (VAF 5%), although equivalent protocols have recently been used to exclude genetic drivers of the precancerous niche^58^.

Chemoprevention of lung cancer has a chequered history. It is a compelling proposition, but there are challenges inherent in balancing efficacy with potential toxicity in individuals who may be at risk but are healthy. Hence, most efforts to date have repurposed drugs such as aspirin and metformin that are safe, cheap and have well understood toxicity profiles^59,60^. We now demonstrate that a clinically approved small molecule kinase inhibitor - capivasertib - is effective at preventing the NTCU-driven phenotype *in vitro* and was not toxic at the effective dose. Further, it was effective at reverting the carcinogen-induced phenotype once established. This is an important proof-of-principle that targeted inhibitors may have the potential to be repurposed for chemoprevention and suggests a potential role in distinct clinical scenarios - high-risk individuals with no evidence of disease and those with biopsy proven established preinvasive disease. However, oral capivasertib has a significant side-effect profile that would preclude its use for this indication.

We therefore modelled the impact of repurposing capivasertib as a nebulised preparation and showed that intermittent therapy prevents the emergence of the carcinogen-associated phenotype at achievable doses and without overt toxicity to normal airway epithelium. Further, nebulised capivasertib reversed the NTCU-phenotype once established. It is therefore plausible that inhalation of capivasertib may be sufficient to rebalance the clonal dynamics within the airway in favour of “normal” clones, and could be infrequent. The inhaled delivery of small molecule targeted inhibitors to the airway is readily achievable and is actively being pursued for airways disease and pulmonary fibrosis^61,62^. Our work therefore raises the intriguing possibility of future inhaled therapeutics for the prevention of lung cancer.

## Materials and Methods

### Patient sample selection

HBECs were obtained either commercially (Lonza, Cat. no. CC-2540 - non-smoking male Donor) or derived from primary airway samples from patients at Cambridge University Hospitals NHS Foundation Trust (CUHFT), with ethical approval (Models of Lung Disease - Research Ethics Committee Reference - 19/SW/0152). Primary cells from CUHFT patients were isolated from bronchial brushings collected from the main airway during bronchoscopy. For patient sample meta data, see **Supp. Table 1**. Patient samples for histology and immunohistochemical analysis were obtained from The Royal Papworth Hospital Research Tissue Bank (Ethical approval 23/EE/0198), reviewed by the East of England - Cambridge East Research Ethics Committee.

### ALI culture of HBECs

ALI cultures were generated as previously reported^63,64^. In brief, HBECs were used at passages up to a maximum of 5. They were expanded from bronchial brushings in PneumaCult™-Ex Plus medium and then detached, suspended in 200μl of supplemented PneumaCult™-Ex Plus medium and seeded on 24-well Transwell inserts (0.4μm pore size; Cat# 353095, Falcon) pre-coated with rat tail type I collagen (Cat# 354236, Corning). After 24 hours, media in both the apical (200µl) and basolateral (500µl) compartments were refreshed. Once confluent (generally Day 2-3), the apical medium was removed to initiate ALI (designated ALI day 0), and 300µl basolateral medium was replaced with HBEC differentiation media (PneumaCult™-ALI, Cat# 05021; Stemcell). Media was replenished every two or three days, and the apical surface washed twice weekly with warm phosphate buffered saline (PBS) to clear accumulated mucus and secretions. These ALI epithelioid cultures were differentiated for a minimum of 28 days prior to experimental use. For cell collection, cultures were washed (PBS) and then treated with Accutase (Stemcell Technologies; Cat. no. 07920) at 37°C for 15 mins. Cells were gently dislodged by pipetting, after which the enzyme activity was quenched using DMEM containing 10% fetal bovine serum. The cell suspension was subsequently centrifuged at 300 × g for five mins at room temperature to pellet cells.

### Capivasertib treatment

For mimicking oral (“systemic) drug delivery (via the blood to the lung) capivasertib was added to the basolateral compartment on a cyclic schedule of four days on and three days off per week. The apical surface was washed with PBS at least once weekly.

For nebulised (“inhaled”) delivery of capivasertib, a prototype version of the commercially available VITROCELL Cloud MAX (VITROCELL Systems, Waldkirch, Germany) for 6-well Transwell insert was employed.

### Histology

Transwell cultures were washed three times with PBS and fixed in 4% paraformaldehyde for 15 mins at room temperature. Membranes were excised from the inserts using a scalpel and paraffin embedded. Sections (4µm) were cut and stained according to standard protocols.

### Immunofluorescence staining

Fully differentiated Transwell cultures were washed three times with PBS and fixed in 4% paraformaldehyde for 15 mins at room temperature before immunostaining. The following antibodies were used: KRT5 antibody (Abcam, ab52635), p63 antibody (Abcam, ab124762), acetylated α-tubulins (Sigma–Aldrich, T7451), VIM antibody (Abcam, ab92547), SOX9 antibody (Cell Signaling Technology, 82630), KRT7 antibody (Abcam, ab181598), KRT17 antibody (Abcam, ab109725), γH2AX antibody (Cell Signaling Technology, 9718), cleaved caspase-3 antibody (Cell Signaling Technology, #9664) and Hoechst dye solution (100µg/ml) for nuclear counterstaining.

### Immunohistochemistry

Formalin-fixed, paraffin-embedded (FFPE) tissue sections (4μm) were mounted on positively-charged slides and baked at 60°C for 1 hour. Sections were deparaffinized in xylene and rehydrated through a graded ethanol series to water. Antigen retrieval was performed using heat-induced epitope retrieval in citrate (pH 6.0) or Tris–EDTA (pH 9.0) buffer at 95–100°C for 10–20 mins.

Endogenous peroxidase activity was quenched with 3% hydrogen peroxide. The following steps were performed using VECTASTAIN® Elite® ABC-HRP Kit (Peroxidase, Universal, PK-6200) according to manufacturer recommendation. Sections were blocked in 5% normal horse serum, then incubated with primary antibodies either for 1 h at room temperature or overnight at 4°C. The following primary antibodies: (KRT5 (Abcam ab52635), SOX9 (Cell Signalling 82630), SERPINB2 (Abcam, ab267463), p40 (Biocare Medical, clone BC28)) were used, followed by incubation with HRP-conjugated secondary antibodies. Signal detection was performed using 3,3′-diaminobenzidine (DAB), and nuclei were counterstained with hematoxylin.

Slides were dehydrated, cleared, and mounted using a permanent mounting medium. Appropriate positive and negative controls were included for all experiments.

### Statistical and computational analyses

All analyses were performed in R version 4.5.0 using packages from the Bioconductor project, release 3.21^66,67^. Figures were generated with ggplot2 (v4.0.2), together with tidyverse (v2.0.0), patchwork (v1.3.0), cowplot (v1.1.3), ggpubr (v0.6.2), ggrepel (v0.9.6) and pheatmap (v1.0.12). With the exception of WGS, all downstream analysis scripts were written as R Markdown documents; code will be available at https://github.com/holly-giles/ntcu_model_2026.

### Sequencing and processing of single cell transcriptomics data

Cell pellets were collected from experimental samples and submitted to the Cancer Research UK (CRUK) Genomics Facility for single cell transcriptomics sequencing. Libraries were prepared using the 10x Genomics Chromium Single Cell 3′ Gene Expression platform according to the manufacturer’s instructions. Sequencing was performed on an Illumina NovaSeq 6000 instrument using an S4 flow cell with XP workflow and a sequencing configuration of up to 200 million reads per library.

Raw sequencing data was processed using the 10x Genomics Cell Ranger (v7.0.1) pipeline for demultiplexing, alignment, and generation of gene expression matrices. Reads were aligned to the human reference genome (10X Human GRCh38 2020 A build). Filtered feature-barcode matrices were used for downstream single cell transcriptomic analysis.

### Quality control, cell-type clustering, and cell-type identification

Single cell analysis was performed using Seurat version 4.3.0.1^68^ unless otherwise stated. Cells expressing fewer than 1000 unique feature counts were excluded from the analysis. Additionally, cells with over 100,000 unique molecular identifiers (UMIs) or mitochondrial content exceeding 17% were discarded. Doublets were filtered with the scDblFinder software^69^: 4-12% of cells were filtered out in each library. This resulted in a final dataset of 41,113 cells for downstream analysis (Control (three samples): 15,165 cells, 3µM NTCU (three samples): 14,680 cells, 10µM NTCU (three samples): 11,268 cells, full sample details in **Supp. Table 1**).

Expression matrices were normalised via the *NormalizeData* function. The *FindVariableFeatures* function was applied to select the top 2000 variable genes, and data from the three patient samples and three conditions was integrated using the *IntegrateData* function.

Next the data was scaled, regressing out variation due to mitochondrial RNA content, followed by principal component analysis (PCA). The first 20 principal components and a resolution of 0.35 were used with the *FindClusters* function to identify 16 clusters. Cell clusters were annotated using a combination of visualising the top eight expressed genes in each cluster, and cross-referencing with the canonical markers of the major cell types (ciliated, secretory, basal, ionocyte, neuroendocrine, tuft, cycling basal, suprabasal, deuterosomal), defined in the literature^9,12,13–17^ (**Supp. Table 2).** Positive markers of each cluster were found using the *FindAllMarkers* function with min.pct = 0.3, logfc.threshold = 0.3, only.pos = TRUE. Clusters assigned to the same cell type were aggregated for subsequent analysis. The consequential cell type assignments were manually reviewed for accuracy and visualised in UMAP plots. Cell-type composition testing was performed using speckle (v1.8.0), comparing 10µM NTCU with Control. .

### Differential expression analysis

In each case, gene-level significance was assessed using Seurat’s *FindAllMarkers* function, using a two-sided Wilcoxon rank-sum test with Bonferroni correction for multiple testing. Genes were ranked based on log fold change. For the 372 ABC marker genes, the following cut offs were applied: min.pct = 0.3, logfc.threshold = 0.3, only.pos = TRUE, all genes had adj. *p*<0.001.

### Trajectory analysis with Monocle3

Trajectory analysis was performed using Monocle3 (v1.4.26)^18^ and Seurat (v5.5.0)^70^. The integrated Seurat object was converted to a Monocle3 *cell_data_set* object using the SeuratWrappers package (v0.4.0)^70^. To ensure all cell types were analysed within a single connected trajectory, all cells were assigned to a single partition. Cluster identities and UMAP coordinates were transferred directly from the Seurat object to preserve consistency with the cell type annotations described above. A principal graph was learned across the UMAP embedding using the *learn_graph* function with default parameters, and cells were ordered in pseudotime using *order_cells* with basal cells specified as the root population, reflecting their identity as the progenitor compartment. The resulting trajectory and pseudotime values were visualised on the UMAP embedding.

### Cell lineage analysis with Slingshot

Trajectory analysis was performed on the ciliated, ABC, basal and cycling basal cells using Slingshot (v2.18.0)^19^ and Seurat (v5.5.0)^70^. Trajectories were inferred using the *slingshot()* function with default parameters on UMAP embeddings, with cell type annotations defining cluster structure and basal cells specified as the starting population, resulting in two inferred lineages. Statistical significance of the shift in pseudotime density distribution between Control and 10µM NTCU samples was calculated using a Kolmogorov-Smirnov test.

### RNA velocity analysis

Spliced and unspliced count matrices were quantified from Cell Ranger BAM files using velocyto^71^, producing one loom file per sample. Loom files were read into R and combined across the nine libraries. The integrated SeuratObject was converted to a SingleCellExperiment object using SingleCellExperiment (v1.30.1) and spliced/unspliced counts added as additional assays. RNA velocity was estimated using the velociraptor R package^71,72^ (v1.18.0), which provides a Bioconductor interface to the scVelo Python library^20^. Velocity was computed using the dynamical model (*scvelo*, mode = "dynamical") on the 2,000 highly variable genes identified during Seurat pre-processing, using the pre-computed PCA embedding. Velocity vectors were projected into UMAP space via the velocity graph for visualisation.

### Heatmaps

For the single cell heatmaps in (Fig. 2H, 2L, 3H) expression values were converted to z-scores of the log-normalised data from the *ScaledData* slot of the Seurat Object. Gene expression was therefore centred and scaled across all cells in the Seurat Object and plotted with pheatmap (v1.0.12). To improve visual contrast and reduce the influence of outliers, scaled values were capped at ± 2.5 before plotting. A random sample of 2000 cells is shown in each case.

### Gene Set Enrichment Analysis

Log_2_FCs for each gene were calculated using the *FindAllMarkers* function. Genes were then ranked according to the resulting log_2_FC statistics and GSEA was performed using the clusterProfiler package (v4.16.0)^73^, with the fgsea algorithm (v1.34.2) and using Hallmark pathway gene sets from the MSigDB database (v25.1.1)^74^.

### Western blotting

Western blotting was performed using standard procedures. KRT5 (Abcam, ab52635), p53 (Santa Cruz Biotechnology, sc-126), total AKT (Cell Signaling Technology, 9272), phospho-AKT (Ser 473) (Cell Signaling Technology, 9271), phospho-AKT (Thr 308) (Cell Signaling Technology, 9275) and β-actin antibody (Santa Cruz Biotechnology, sc-69879) were used.

### Preparation of DNA for Whole Genome Sequencing (WGS)

Genomic DNA was extracted from differentiated ALI cultures exposed to either vehicle control or NTCU (10µM) for six weeks using the DNEasy Tissue and Blood Kit (Qiagen, Cat no. 69504).

### Preparation of DNA for 46-gene panel

The genomic DNA was extracted using the QIAamp DNA Micro Kit (Cat no. 56304) according to the manufacturer’s recommendations before quality control and sequencing by Novoseq.

### Processing and analysis of 46-gene panel DNA data

DNA was analysed using the Oncomine Precision Assay X on the Ion Torrent Genexus Integrated Sequencer (Thermo Fisher Scientific). The Integrated Sequencer automated library preparation, templating, sequencing, variant calling and determination of heterozygosity according to the manufacturer’s protocol. Reads were aligned to the reference human genome hg19 (GRCh37). The assay interrogates 250 mutational hotspot regions across 46 cancer-associated genes for single nucleotide variants (SNVs) (**Supp. Table 4**). Variants were called and classified as PRESENT, NO CALL or ABSENT using the Oncomine Reporter with default manufacturer filters. The reported validated limit of detection for this assay is a VAF of 5% at a minimum coverage of 250 reads.

The assay detected 20 present variants in each sample, with a VAF of >5%. Somatic variants were defined as those detected in NTCU-treated samples but absent in matched vehicle controls (**Figures 4A, 4B**). Each variant was observed in both control and NTCU samples and had a VAF of 29–100%, consistent with germline polymorphisms. The Oncomine report confirmed that all 20 detected variants represented common germline polymorphisms (population Minor Allele Frequency >1%) or intronic/non-coding variants, with no variants of clinical significance identified.

Two additional variants above the 5% VAF threshold (GNA11 p.Q209R and p.Q209K) were excluded from the analysis: both were present in NTCU and control samples at similar frequencies and flagged as NO CALL by the Ion Torrent variant caller due to extreme strand bias (score >0.99).

### Sequencing and alignment of WGS data

Genomic DNA from HBEC samples was randomly sheared, end-repaired, A-tailed, and ligated with Illumina adapters. Adapter-ligated fragments were size-selected, PCR-amplified, and purified. Libraries were quantified by Qubit and qPCR with size distribution assessed on a fragment analyzer, then pooled and sequenced on an Illumina NovaSeq X Plus (PE150). Raw reads were filtered to remove adapter-containing reads, reads with >10% ambiguous bases, and reads with >50% low-quality bases (Q ≤ 5). Clean reads showed Q30 > 92% and effective rates > 99% across all lanes.

### Processing of WGS data

Somatic mutation calling was performed as described previously^75^, comparing differentiated ALI cultures exposed to NTCU (10µM) with vehicle control as the reference. Briefly, the mutation calling pipeline was containerised within dockstore-cgpwgs (https://github.com/cancerit/dockstore-cgpmap) (v2.1.1), implementing Caveman^76^ (Cancer Variants through Expectation Maximisation, v1.13.15) for somatic substitution calling, Pindel^77^ (v3.2.0) for detection of indels, and BRASS (BReakpoint AnalySiS, v6.2.1) for structural rearrangements. ASCAT^78^ (v4.2.1) was used for copy number analysis. Additional filtering was applied as follows: SBSs were filtered based on a PASS, CLPM = 0.00, and ASMD > = 140, small indels were filtered for QUAL > = 250 & REP < 10, and structural variants filtered for PASS and BRASS assembly score >0 indicating successful *de novo* local assembly of the breakpoint junction sequence using Velvet.

### Analysis of ABC marker gene expression in clinical bronchial biopsy specimens

Gene expression data were obtained from GEO (accession GSE33479, Agilent GPL6480, Whole Human Genome 4x44K two-colour array)^35^, comprising microarray profiles of 122 bronchial biopsies from 77 patients (35 former, 42 current smokers) across eight histological stages: normal epithelium, hyperplasia, metaplasia, mild dysplasia, moderate dysplasia, severe dysplasia, carcinoma in situ, and invasive squamous cell carcinoma, as graded by independent pathological review. In this dataset, each biopsy (Cy5) was hybridised against a common pooled reference RNA from normal biopsies of 16 never-smokers (Cy3). Each measurement therefore reflects relative transcript abundance: the Cy5/Cy3 intensity ratio for each probe, comparing the biopsy of interest to the pooled never-smoker normal reference.

Data were accessed using the GEOquery package^79^, ratios were already normalised as described in the manuscript^35^, and no further preprocessing was applied.

An ABC gene signature was derived from the top 20 marker genes identified by *FindAllMarkers* in the NTCU single cell transcriptomics dataset (described above). Probes were mapped to gene symbols using the feature annotation provided with the dataset, and probes lacking an assigned gene symbol were excluded. Where multiple probes mapped to the same gene, expression values were collapsed to a single gene-level estimate by taking the mean across probes, using the *avereps* function from the limma package^79,80^. Genes in the ABC signature were intersected with genes present on the array, and the resulting matched gene set of 20 genes was used for downstream analyses.

For each sample, a composite ABC signature score was calculated as the mean expression across all matched ABC marker genes. Six driver genes (*KRT7, MACC1, ANXA3, SERPINB2, SFN, KRT23*) were examined individually. Trends with stage were tested by Spearman correlation.

To test specificity, we generated 20 control gene sets (20 genes each, sampled without replacement) from all single cell transcriptomics cluster marker genes (excluding the ABC top-20), restricted to genes detectably expressed (mean > 1) in the Mascaux array. Scores were computed identically and compared across stages.

### LDH cytotoxicity assay

LDH assay was conducted as described previously. For positive and negative controls, 50μL of 10× lysis buffer or ultrapure water, respectively, was applied to the apical surface of HBEC-ALI or nasal ALI Transwells and incubated at 37°C for 45 mins. The apical chamber was then washed using 500μL of basal medium. For experimental conditions, the apical surface was similarly washed with 500μL of basal medium. All conditions were prepared in triplicate. LDH release in each sample was quantified using the CyQUANT LDH Cytotoxicity Assay Kit (C20300; Invitrogen) according to the manufacturer’s instructions. Briefly, 50μL of the collected sample was mixed with 50μL of reaction mixture and incubated at room temperature for 30 mins. The reaction was then stopped, and absorbance was measured at 490nm and 680nm.

Cytotoxicity (%) was calculated using the following formula:

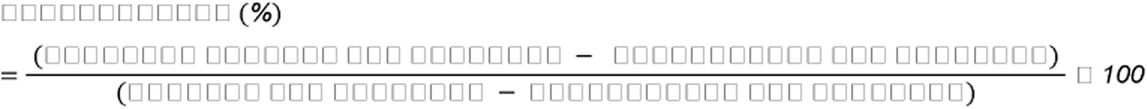

### TEER

TEER was performed as previously described^81^ using EOVM3 (World Precision Instruments). Before measurement, plates were equilibrated to room temperature for 15–30 mins. Resistance was measured using an epithelial voltohmmeter with sterilised electrodes placed in the apical and basolateral compartments. Blank values from cell-free inserts were subtracted. Final TEER values were calculated by multiplying resistance by membrane area and expressed as Ωcm².

### Use of AI tools

Anthropic’s Claude (Claude Code and Claude large language models) was used to assist in writing and debugging the R analysis and figure-generation code used in this study, and to help draft portions of the Methods. All code and text were reviewed, tested, and edited by the authors, who take full responsibility for the accuracy and integrity of the work. No AI tool was used to interpret experimental data.

### Code availability

All code to reproduce the single cell transcriptomics, 46-gene panel and clinical sample transcriptomics analyses is available for reviewers - it has been uploaded as supplemental info (and will be available on GitHub). For the WGS analysis, the mutation calling pipeline was containerised within dockstore-cgpwgs (https://github.com/cancerit/dockstore-cgpmap) (v2.1.1).

### Data sharing statement

Processed data objects for the single cell transcriptomics, 46-gene panel, WGS and clinical sample transcriptomics analyses performed in this study will be available on GitHub/Online. For all original sequencing data, we will deposit in the European Genome-phenome Archive.

### Analysed data from previous studies

We obtained microarray profiles of bronchial biopsies from the NCBI GEO database (Mascaux et al. 2019, GSE33479)^35^. We obtained marker gene lists and associated logFC values for the aberrant basaloid, transitional and Hillock-like populations described by Adams et al 2020^31^, Habermann et al. 2020^31,32^, Strunz et al. 2020^33^, Lin et al. 2024^34^ and Gomez-Lopez et al. 2025^9^ from the published marker tables of these studies.

## Funding

F.M. and W.G. were supported by the BBSRC (BB/W014564/1) and the NC3Rs (NC/S001204/1). This research was supported by the NIHR Cambridge Biomedical Research Centre (NIHR203312). The views expressed are those of the authors and not necessarily those of the NIHR or the Department of Health and Social Care

Strategic partnership between University of Cambridge and Ludwig-Maximilians-Universität München (LMU)

National Research Foundation of Korea grant funded by the Ministry of Science and ICT (RS-2025-18362970)

Brain Pool Plus Fellowship Program funded by the Ministry of Science and ICT (RS-2025-25427881)

We donors and the Royal Papworth Hospital Research Tissue Bank for the collection, storage, and provision of human biological materials/surgical samples. We thank the CRUK-Cambridge Institute sequencing facility, Jeffrey Cheah Building IT Services and members of the Han Lab for advice on statistics and analysis. H.A.R.G. thanks Christ’s College for supporting her as a Bye-Fellow during this project.

## Author contributions

F.M. and W.G. conceptualised the study. F.M., H.A.R.G., O.S., S.N-Z., L.D., M.G. and N.H. conceptualised the data analysis.

F.M., W.G., O.S., A.H., A.O.Y., H.A.R.G., H.D., S.N-Z. and N.H. designed the experiments.

W.G., A.C. and C.G. performed the experiments.

H.A.R.G. performed data processing and curation, developed and maintained the analysis code, and wrote the software documentation.

J.X. and H.A.R.G. ran the Slingshot analysis.

Y.M, D.B. and H.D. ran the WGS analysis, H.A.R.G. ran the 46 gene panel analysis.

L.D. and M.G. analysed the histology.

W.G., H.A.R.G., H.D., S.N-Z, F.M. and N.H interpreted the analyses.

H.A.R.G. , W.G. produced the figures (with support from F.M. and N.H).

A.Ö.Y., O.S. supervised aspects of the drug delivery experiments

S.N-Z. supervised the whole genome sequencing

F.M., H.A.R.G., W.G., S. N-Z., O.S. and N.H. drafted the manuscript. All authors reviewed and edited the manuscript.

W.G. and H.A.R.G. contributed equally to this work. F.M. and N.H. jointly supervised this work.

## Conflicts of interest

N.H. is the co-founder and Chief Technology Officer of CardiaTec Bio, a company developing therapeutics for cardiovascular diseases, and the co-founder of KURE.ai, which focuses on AI-driven oncology drug discovery. N.H. also serves on the Scientific Advisory Board of the Institute of Cancer Research (ICR). These affiliations are unrelated to the subject matter of this manuscript. The other authors declare no competing interests. FM has had research funding in kind from AstraZeneca through the provision of materials in the past. He currently co-supervises a PhD student in an unrelated project with an AstraZeneca employee.

