## Supplemental Figure and Table Legends for "Carcinogen-induced preinvasive human squamous lung cancer is characterised by a novel atypical basal cell state and presents opportunities for chemoprevention"

### **Supplementary Figure and Table Legends**

##### **Supp. Figure 1.**

**Supporting single cell transcriptomics data for cell type annotation, proportions, pseudotime and RNA velocity.**

**(A)** Dot plot showing expression of canonical marker genes across all nine annotated cell types. Dot size indicates the proportion of cells expressing each gene; colour indicates mean scaled expression.

**(B)** Stacked bar chart of cell type proportions per sample, grouped by patient sample (1-3) and N-nitroso-tri-chloroethylurea (NTCU) treatment dose (Control, 3µM, 10µM).

**(C)** Density plot of Slingshot pseudotime values for Lineage 2, split by treatment group.

**(D)** RNA velocity Uniform Manifold Approximation and Projection (UMAP) overlaid with velocity vectors (arrows), coloured by cell type, showing directional transcriptional dynamics across the epithelial hierarchy.

**(E)** Violin plot of *VIM* log-normalised expression across all cell types.

**(F)** Gene Set Enrichment Analysis (GSEA) summary table showing the top five enriched and top five depleted Hallmark pathways in basal cells from NTCU-treated versus control cultures. NES = Normalised enrichment score, pathways ranked by p value, p values adjusted by Benjamini-Hochberg FDR.

##### **Supp. Figure 2.**

**Supporting data for ABC marker genes, GSEA and DNA damage response.**

**(A)** Violin plots showing *SOX2* and *SOX9* log-normalised expression in atypical (ABC) and basal cells, split by NTCU treatment dose (Control, 3µM, 10µM). *SOX9* expression is upregulated and SOX2 downregulated in ABCs in 10µM versus Control (both adj. *p*<0.001, Wilcoxon rank-sum, Bonferroni corrected).

**(B)** Feature plots showing *KRT7* and *KRT17* expression projected onto the full UMAP of all cells. *KRT7* and *KRT17* are both markers of ABCs (both adj. *p*<0.001, Wilcoxon rank-sum, Bonferroni corrected).

**(C)** Immunofluorescence (IF) image of Control and NTCU-treated ALI cultures, stained for acetylated tubulin (green), KRT17 (red) and nuclei (blue). Image is representative of three replicates.

**(D)** Violin plot of *KRT17* log-normalised expression in atypical (ABC) and basal cells, split by treatment. KRT17 expression increased in ABC population in response to NTCU (adj. *p*<0.001, Wilcoxon rank-sum, Bonferroni corrected).

**(E)** GSEA summary table showing the top five enriched and top five depleted Hallmark pathways in atypical cells (ABCs) compared to all other cell types. Gene set enrichment analysis (GSEA) performed on ranked log_2_FC values, using the fgsea algorithm. DNA Repair, Benjamini-Hochberg adj. *p*=0.028).

**(F)** Violin plot of *H2AFX* log-normalised expression in atypical (ABC) and basal cells, split by treatment.

**(G)** Violin plot of *TP53* log-normalised expression in atypical (ABC) and basal cells, split by treatment.

##### **Supp. Figure 3.**

**The atypical cell (ABC) population compared to previously described aberrant basaloid and hillock-like epithelial states.**

Heatmaps of the top 50 marker genes of five published epithelial cell states showing expression across the nine cell types we identified in the NTCU single cell transcriptomics dataset. **(A–C)** Signatures derived from human studies:

**(A)** aberrant basaloid cells from idiopathic pulmonary fibrosis (IPF) and chronic obstructive pulmonary disease (COPD) lung (Adams et al. 2020; 49/50 genes detected),

**(B)** KRT5^-^/KRT17^+^ epithelial cells from IPF lung (Habermann et al. 2020; 49/50 genes detected)

**(C)** KRT4/KRT13-high intermediate cells associated with tobacco exposure (Gómez-López et al. 2025; 50/50 genes detected).

**(D, E)** Signatures derived from mouse studies, mapped to human orthologues:

**(D)** KRT8^+^ alveolar differentiation intermediate (ADI) cells arising after bleomycin injury in mice (Strunz et al. 2020; 50/50 genes detected) and

**(E)** hillock basal cells of the normal airway (Lin et al. 2024; 49/50 genes detected).

For each signature, marker genes were taken from the published marker table of the original study and ranked by the reported log fold change. Data from Hoffman et al. 2025 is not shown as the marker table was not available (comparison with ABCs was therefore based on selected quoted markers from text). Values are mean log-normalised expression per cell type, pooled across all three patient samples and all NTCU doses, z-scored by gene (red, high; blue, low). Rows are ordered in descending order, according to reported log fold change of the marker genes. ABC population is highlighted in green.

##### **Supp. Figure 4.**

**Whole-genome sequencing for two patient samples comparing two treatments: 10µM NTCU vs Control.**

**(A)** Single nucleotide variant (SNV) mutational profile for Patient sample 1 showing trinucleotide context counts across the 96 SNV mutation classes.

**(B)** SNV mutational profile for Patient sample 3.

**(C)** Mutational profile for small insertions and deletions (indels) for Patient sample 1, showing counts by indel type.

**(D)** Indel mutational profile for Patient sample 3.

Profiles are shown for NTCU-treated samples relative to matched controls.

##### **Supp. Figure 5.**

**Random gene set controls for clinical validation analysis.**

**(A)** Median atypical (ABC) signature score (red) compared to the median values for 20 randomly sampled gene sets of equivalent size (n = 20 genes each, grey) across histological stages in the Mascaux et al. dataset. Random gene sets are drawn from the full pool of single cell transcriptomics cluster marker genes i.e. all genes are epithelial-related marker genes in our dataset. SCC = Squamous cell carcinoma.

(B) Serial sections from three independent metaplastic foci in Patient C were stained with hematoxylin and eosin (H&E) for KRT5, p40, and SERPINB2 (green, with Hoechst nuclear counterstain). Squamous metaplasia strongly expresses KRT5, p40 and SERPINB2. Scale bars, 300μm.

##### **Supp. Figure 6.**

**Target engagement and extended histology data for capivasertib treatment.**

**(A)** Western blot showing levels of total AKT and phosphorylated AKT in Air-Liquid Interface (ALI) cultures treated with and without NTCU and basal capivasertib treatment. Human bronchial epithelial cells **(**HBECs) cultured for 48 hours.

**(B)** Representative H&E-stained sections following nebulised capivasertib treatment in the presence of NTCU. Scale bar, 300μm. Representative histology from experiments on two patient samples using a basal dose equivalent to 3µM.

#### **Supp. Table Descriptions**

##### **Supp, Table 1.**

*Metadata for the single cell transcriptomics patient samples.* Table includes details of the nine libraries (three patient samples 1 - 3, each treated with vehicle control and NTCU at 3µM and 10µM) alongside patient sample characteristics, treatment conditions and post-quality-control cell numbers (41,113 cells total). The same patient samples 1 -3 were used for the genomics analysis. .

##### **Supp. Table 2.**

*Canonical marker genes used for cell-type annotation.* Marker genes for each airway epithelial cell type (basal, cycling basal, suprabasal, ciliated, deuterosomal, secretory, ionocyte, neuroendocrine, tuft), were compiled from published datasets and used to annotate the single-cell clusters.

##### **Supp. Table 3.**

*Marker genes of the atypical basal cell (ABC) population.* The 372 genes significantly upregulated in the ABC cluster relative to all other cells (Seurat FindAllMarkers; two-sided Wilcoxon rank-sum test, only.pos = TRUE, min.pct = 0.3, logfc.threshold = 0.3), ranked by average log2 fold change. All genes have Bonferroni-adjusted P < 0.05.

##### **Supp. Table 4.**

*Details of genes and amplicons in the targeted deep-sequencing panel.* The 46 cancer-associated genes (250 mutational hotspot regions) interrogated by the Oncomine Precision Assay to screen control and NTCU-exposed cultures for somatic variants.
